# Engineering 3D Vascularized Adipose Tissue Construct using a Decellularized Lung Matrix

**DOI:** 10.1101/2021.06.22.449445

**Authors:** Megan K. DeBari, Wai Hoe Ng, Mallory D. Griffin, Lauren E. Kokai, Kacey G. Marra, J. Peter Rubin, Xi Ren, Rosalyn D. Abbott

**Author notes:** Co-first author (Megan K. DeBari, Wai Hoe Ng).

## Abstract

Critically sized defects in subcutaneous white adipose tissue result in extensive disfigurement and dysfunction and remain a reconstructive challenge for surgeons; as larger defect sizes are correlated with higher rates of complications and failure due to insufficient vascularization following implantation. Our study demonstrates for the first-time a method to engineer perfusable, pre-vascularized, high-density adipose grafts that combine patient-derived adipose cells with a decellularized lung matrix (DLM). The lung is one of the most vascularized organs with high flow, low resistance, and a large blood-alveolar interface separated by a thin basement membrane. For our work, the large volume capacity within the alveolar compartment was repurposed for high-density adipose cell filling, while the acellular vascular bed provided efficient graft perfusion throughout. Both adipocytes and hASCs were successfully delivered and remained in the alveolar space even after weeks of culture. While adipose derived cells maintained their morphology and functionality in both static and perfusion DLM cultures, perfusion culture offered enhanced outcomes over static culture. Furthermore, we demonstrate that endothelial cells seamlessly integrate into the acellular vascular tree of the DLM with adipocytes. These results support that the DLM is a unique platform for creating vascularized adipose tissue grafts for large defect filling.

## 1. Introduction

Adipose tissue is a highly vascularized, metabolically active connective tissue that plays a crucial role in supporting normal human appearance, protecting organs, modulating pressure, providing insulation, and functioning as an endocrine regulator. The appearance and function of subcutaneous adipose tissue can be compromised by traumatic injury (i.e. motor vehicle accidents, burns), ablative procedures (i.e. mastectomy), or congenital malformations (Romberg’s disease, Poland syndrome) leading to contour abnormalities and exposure of vital organs. There is a strong clinical need for adipose tissue replacements, as pathological changes and defects of the subcutaneous adipose tissue cause extensive disfigurement and dysfunction and remain a reconstructive challenge for surgeons, where increasing defect size is correlated with an increase in complications and failure [1, 2].

Currently large subcutaneous defects are treated with autologous adipose grafts which result in donor-site morbidity and have limited applicability to patients with minimal body fat. For lean patients, the most viable option is allotransplantation which poses a significant risk of allograft rejection and requires lifelong immunosuppression therapy.[3] Tissue engineered adipose grafts using patient-derived cells hold promise in resolving these limitations. However, it remains challenging to engineer large-volume (>200 cm^3^) adipose grafts due to insufficient or delayed vascularization following implantation, which leads to a rapid loss of cell viability, compromised host integration, and limited long-term stability [4-7]. Innovation is needed to engineer critically sized adipose tissue implants that: (1) can fill large tissue defects; results in minimal donor site morbidity and can be implemented in lean patients; and (2) eliminates the need of immunosuppressive therapy and it’s wide variety of side effects [8].

Our overall objective was to develop a biomaterial strategy to enable the engineering of perfusable, pre-vascularized, high-density adipose grafts using patient-derived adipose cells. To achieve this, our approach uses the decellularized lung matrix (DLM) as a unique biomaterial scaffold. The lung is one of the most vascularized organs with high flow, low resistance, and a large blood-alveolar interface separated by a thin basement membrane (<1µm [9]). Accordingly, the DLM offers the following unique advantages: (1) its large volume capacity within the alveolar compartment can be repurposed for high-density adipose cell filling; (2) it offers a preserved, acellular vascular bed allowing efficient graft perfusion and pre-vascularization; (3) the alveolar compartment shields buoyant and fragile adipocytes [10, 11] from high shear stresses that compromise their viability [12] by confining high flow rates to the vascular bed where it is required for proper endothelial functionality;[13] (4) it will not elicit an immune response by allogeneic or xenogeneic recipients;[14-21] and (5) the DLM is scalable, and can be manufactured from rat, pig, and donor human lungs to meet the diverse size requirements for filling different defects.[22-26] In the following studies we use rat as our DLM as a proof of concept, with the long-term goal of scaling up in future studies to larger animal scaffolds.

## 2. Results & Discussion

Prior to using the decellularized lung matrix (DLM), it was important for us to establish that all processing steps required to homogenize, breakup, and filter the adipose cells did not negatively affect the cellular viability, metabolism (Resazurin), and functionality (glycerol secretion). To do this, we used our already established silk scaffold system for adipose tissue culture that is straightforward to setup and evaluate [27-30]. Once we verified the tissue processing did not affect cellular functions and viability, we cultured adipocytes and human adipose derived stem cells (hASCs) in the DLM scaffold in static culture to ensure interactions with the DLM did not affect their morphology and function (in the absence of perfusion). Next, we sought to compare static and perfusion culture for maintaining metabolism (Resazurin) and morphology in culture. Once we determined perfusion culture resulted in enhanced metabolism over static conditions, we proceeded to use the DLM perfusion culture system to seed endothelial cells in co-culture with adipocytes towards a proof of concept fully vascularized adipose tissue construct.

### 2.1. Adipocyte Isolation

In our initial attempts to seed adipocytes into the DLM via the tracheal outlet, we found that unfiltered, minimally processed adipose tissue caused excessive blockage of airway passages and ruptured the DLM (data not shown). Therefore, to enable effective adipocyte loading into the entire alveolar space, we explored different adipocyte isolation methods to mitigate clogging in the bronchioles. Clinically, adipose tissue is harvested using a cannula and then mechanically disrupted with commercially available devices. In order to assess whether our approach would be translatable to clinically used techniques we sought to compare our traditional bench-top methods of adipose tissue processing with a blender to a clinically relevant cannula. Following tissue extraction, we further disrupted the tissue with a LIPOGEMS kit (Figure 1 A). LIPOGEMS is an FDA cleared medical technology that homogenizes adipose tissue for injection into injured or damaged areas to support repair and healing [31]. Although the LIPOGEMS kit does aid in extracellular matrix (ECM) breakdown, there were occasionally large ECM fragments that persisted. To remove these large fragments, a secondary filter (1 mm) was introduced. This resulted in four experimental groups (that were all processed by the LIPOGEMS kit): Cannula, Cannula+Filter, Blender, and Blender+Filter (Figure 1 A).

**Figure 1.**
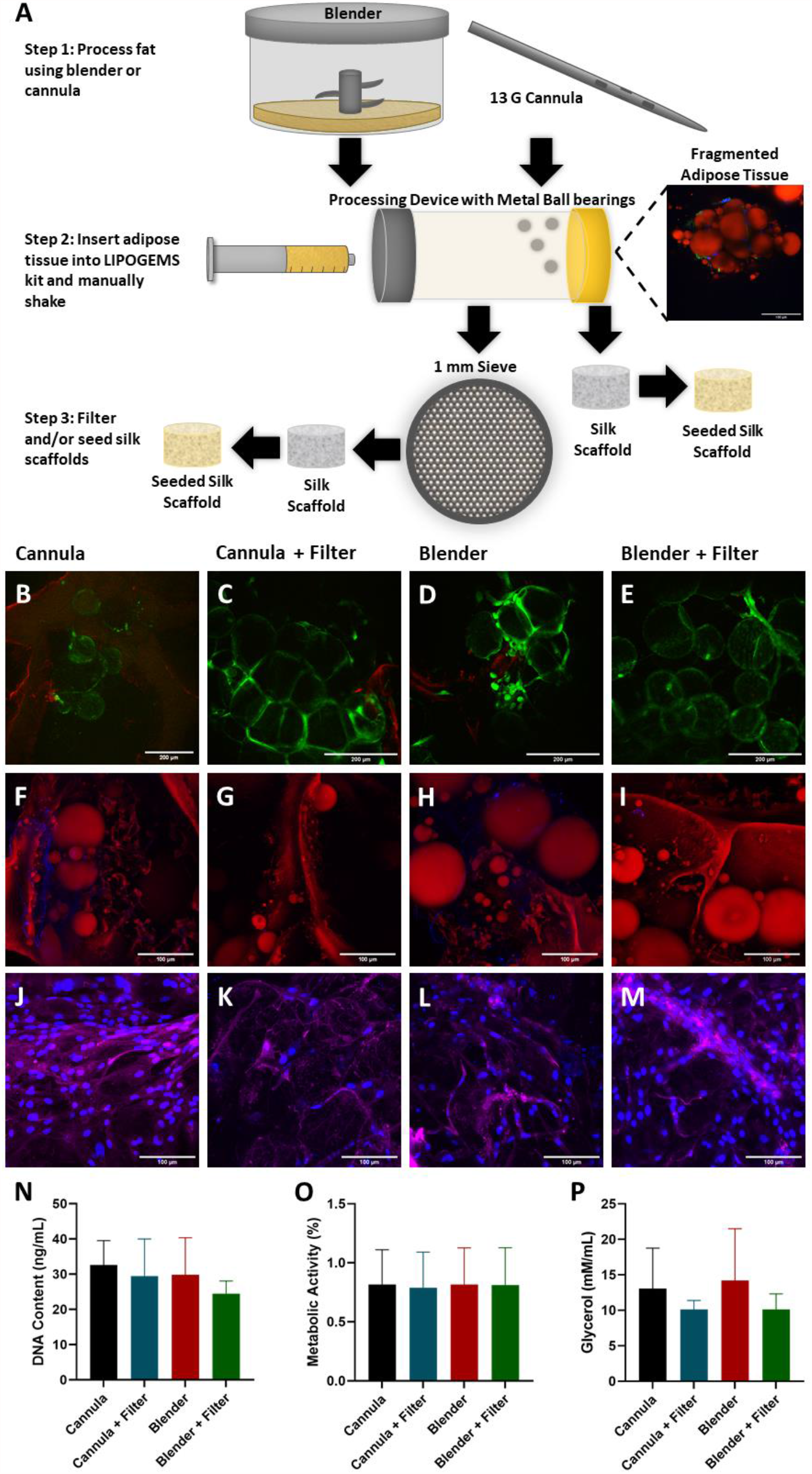
There were no significant differences in adipocyte viability, morphology, or function between the examined tissue processing techniques (blender+LIPOGEMS, cannula+LIPOGEMS, and additional filter). (A) Schematic showing adipose tissue processing steps for initial isolation experiments. Whole adipose tissue was processed to break up ECM components so the tissue could be perfused into the DLM without causing blockages. Image showing fragmented adipose tissue (red – adipocytes, green – actin, blue – nuclei) was captured after processing by LIPOGEMS kit. Scale bar = 100 µm. (B-E) After 4 days of culture, live/dead staining (green – live cells, red – dead cells/silk scaffold) indicated that all isolation processes resulted in high cell viability. Scale bars = 200 µm. (F-I) Second harmonic generation (SHG) was used to image collagen deposits (red – adipocytes/ silk scaffolds, blue – collagen). Tissues that were not filtered through a secondary 1 mm filter had higher amounts of collagen deposits. Scaffolds were cultured for 4 days prior to imaging. Scale bars = 100 µm. (J-M) Scaffolds were imaged to visualize *de novo* protein deposits between day 4 and 7 of culture (blue – Nuclei, purple – de *novo* ECM). All isolation processes resulted in new ECM production indicating that not only were the cells viable, but also capable of remodeling. Scaffolds were cultured for 7 days prior to imaging. Scale bars = 100 µm. (N-P) DNA content, metabolic activity, and glycerol secretions after 4 days of culture. Results show no statistically significant differences between all isolation techniques. Experiment was done in duplicate.

To test our processing methods in a straightforward well-established 3D environment for adipocytes [27-30], we seeded processed adipose tissues in silk scaffolds. While silk scaffolds offer great promise for adipose tissue engineering approaches [27-30, 32-37], due to diffusion constraints and the risk of developing necrotic cores, silk biomaterials alone cannot be used to fill critically sized defects without vascularization (or pseudo vascularization in the form of conduits, perfusion systems, etc.). The DLM is well-suited to address this deficit as it offers a preserved, acellular vascular bed allowing efficient graft perfusion and pre-vascularization [38].

It was critical that not only were adipocytes isolated with minimal ECM to avoid obstructions during seeding, but also that they maintained their viability, morphology, and functionality. After 4 days of culture, live dead staining indicated there was similar adipocytes and stem cells viability in all four experimental groups (Figure 1 B-E), indicating that none of the isolation methods negatively affected cell viability. Next, we sought to determine if the processing steps affected morphology and functionality. A hallmark of a mature adipocyte is the presence of a single unilocular, triglyceride-filled intracellular lipid droplet [39]. Adipocytes in all four experimental groups had the characteristic unilocular morphology (Figure 1 F-I), however, the collagen content in the experimental groups varied. Areas of high collagen content were evident in the groups that were not passed through the secondary filter. As ECM removal is critical to ensure successful seeding into the DLM without rupturing, this led to the conclusion that the secondary filter was a necessary extra step during adipocyte processing prior to DLM seeding. Furthermore, ECM production and remodeling is an essential aspect of a healthy, regenerative tissue [40]. To test that this functionality was maintained, several scaffolds were cultured for 3 additional days and stained to image *de novo* protein synthesis (Figure 1 J-M). All experimental groups had a significant amount of new ECM indicating that cellular functionality was not affected by the extraction, homogenization, and filtering methods.

DNA content, metabolism, and glycerol secretion were all measured after 4 days of culture (Figure 1 N-P). There were no statistical differences between any of the experimental groups with regard to DNA content, metabolism, or glycerol secretion. Although DNA content for the Blender+Filter group appears lower than the other groups, the standard deviation was smaller than the other groups, indicating this isolation method is likely more reproducible than extracting with a cannula. Even with lower DNA content, the Blender+Filter group had similar metabolic activity compared to the other experimental groups. Glycerol secretion was also not significantly different between the experimental groups. However, when the secondary filter was used glycerol secretion appears to be slightly lower with a lower standard deviation. Since the presence of ECM has been shown to affect lipolysis rates [41], the higher ECM content in the Cannula and Blender experimental groups could have contributed to higher levels of glycerol secretion. Overall, these results revealed there was no significant differences between the function, morphology, and viability of the cells isolated using any processing method. Therefore, in subsequent studies the blender was used for adipocyte isolation instead of the cannula because it was not significantly different but could be performed with smaller tissues in a laboratory setting. Due to the lower ECM content when using the secondary filter, lowering the risk of DLM clogging, the filter was also used in all subsequent studies.

### 2.2. Static Culture of Whole Lobes

The DLM is enriched for key ECM components critical for adipogenesis, including collagen I, IV, and VI, as well as laminins and fibronectin [42, 43]. While we speculated that the DLM would be well-suited for supporting the adipose niche, we first sought to evaluate the interaction of adipocytes to the matrix alone without other confounding factors (i.e. no perfusion). To investigate if the DLM could serve as a scaffold framework for supporting adipocytes, the DLM was seeded with fresh adipose tissue and cultured statically for one week (Figure 2 A). One of the major advantages of using the DLM for this study is that different lobes of the DLM could be isolated separately. Hence, individual lobes were harvested on Day 0, 1, 3, and 7 for evaluating morphology (Figure 2 B-E). At all timepoints, images showed healthy looking adipocytes as indicated by spherical unilocular lipid droplets [39]. Intriguingly, we noticed that the stem cells were organized around the adipocytes, recapitulating what is observed *in situ* [44]. Extracellular lipid droplets were also present at each timepoint. Because extracellular lipid droplets were present at Day 0, the extracellular lipid droplets were likely present in the adipocyte solution prior to seeding. As static culture does not encourage diffusion throughout the DLM, lipid droplets did not diffuse into the vasculature as they would *in vivo*. These lipid droplets are characteristic of unhealthy adipose tissue [45] and could leadto ectopic lipid droplets and metabolic dysfunction [46], therefore, we concluded that media perfusion may be beneficial to remove these residual extracellular lipid droplets. Glycerol secretions were measured on day 1, 3 and 7 for the lobe harvested on day 7 (left lobe) (Figure 2 F), where a strong linear relationship was measured between time and glycerol secretion (linear regression R^2^ = 0.9998). This indicates that glycerol levels remained steady throughout the 7-day culture. Overall, the appearance of the adipocytes and stem cells at all timepoints and the steady glycerol levels show that the DLM can support adipocyte morphology and function.

**Figure 2.**
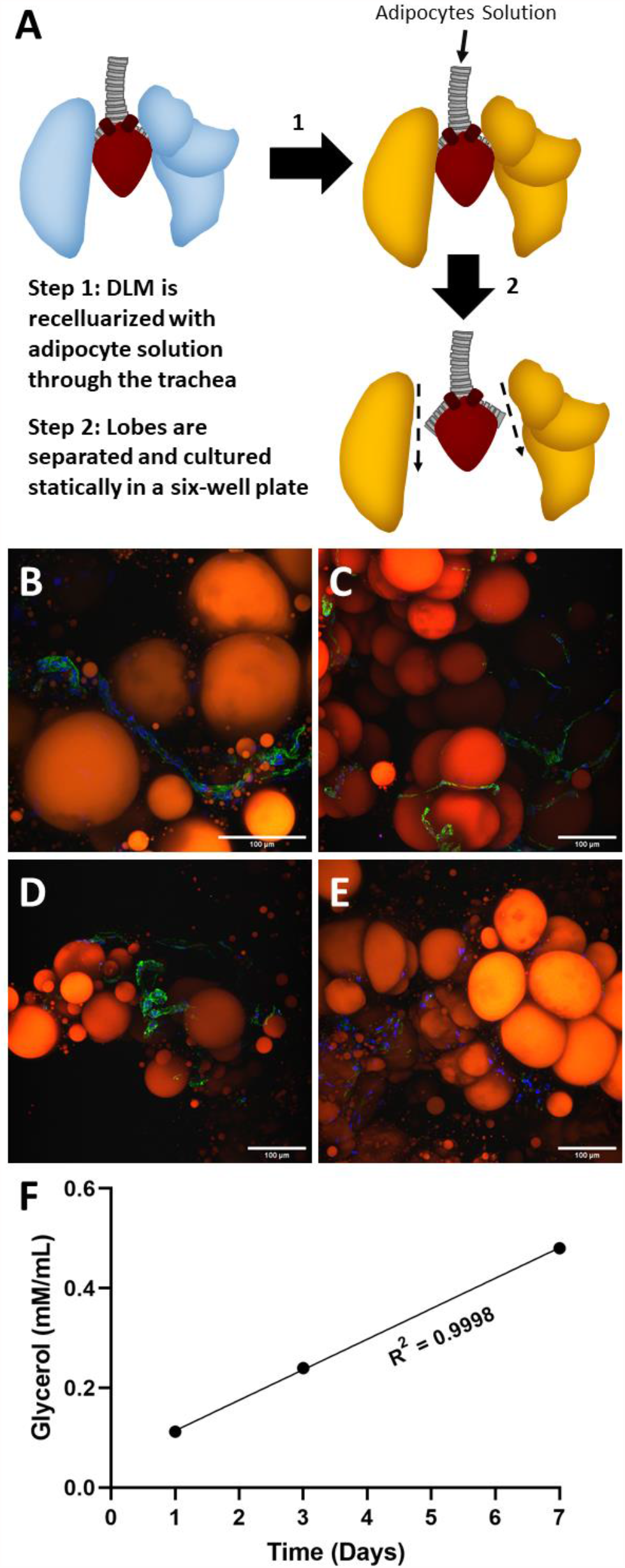
DLM cultured in static conditions supports adipocyte morphology and function. (A) Schematic showing how the DLM was seeded, separated, and cultured statically. Lobes were fixed and stained on days (B) 0, (C) 1, (D) 3, and (E) 7 (red – adipocytes, green – actin, blue – nuclei). Throughout culture adipocytes maintained similar sizes and shapes (unilocular intracellular lipid droplets) in the presence of stromal vascular fraction (SVF) cells (actin positive cells). Extracellular lipid droplets (lipid droplets without nuclei) were present at every timepoint. Scale bars = 100 µm. (F) Glycerol levels for the lobe cultured for 7 days show consistent levels of glycerol production throughout the 7 days indicating the adipocytes remained functional after seeding and throughout culture (linear regression with a goodness of fit = 0.9998).

### 2.3. Culture and Differentiation of Adipose Derived Stem Cells in the DLM

After verifying the DLM could support mature adipocytes, we next wanted to determine whether it could support differentiation of human adipose derived stem cells (hASCs). Mature adipocytes can be readily obtained in large quantities from patients with excess adipose tissue and represents a convenient cell source for adipose tissue engineering as they do not require further expansion or differentiation before cell seeding. On the other hand, hASCs are an attractive option for lean or adolescent patients with minimal body fat as the highly proliferative hASC population can be extracted from a small volume of lipoaspirate and expanded into a large number in culture. To investigate if the DLM supported stem cell differentiation into adipocytes, hASCs were seeded into the DLM, which was then sliced and cultured statically for 4 weeks in either growth or adipogenic differentiation media (Figure 3 A). After 4 weeks, hASCs cultured in differentiation media accumulated intracellular lipid droplets with a mixture of multilocular and unilocular cells (Figure 3 C), while the slices cultured in growth media showed minimal differentiation (Figure 3 B). Not surprisingly, the growth media promoted a small amount of spontaneous lipid accumulation, which we attribute to the low passage number of the hASCs used [47] and the high confluence [48, 49]. Metabolic activity was also assessed in the tissue slices at the end of week 1 and week 4 (Figure 3 D-E). There was no significant difference in the metabolic activity between the growth and differentiation media groups after 1 week of culture. However, after 4 weeks of culture there was a significant difference in metabolic activity between the two groups. This data was normalized to DNA content to account for cell proliferation over the 4 weeks (Figure 3 F). There was a significant difference in DNA content between the two experimental groups, with the slices cultured in differentiation media having a higher DNA concentration. This could have been due to several compounding factors. Endothelial cell viability (from the stromal vascular fraction) could have been negatively affected by the non-differentiating hASCs. Literature has shown that after two weeks of culture endothelial cell viability decreases by more than 20% in the presence of non-differentiating hASCs [50]. Additionally, growth media has shown to result in a slightly lower proliferation rates compared to differentiation media [51]. Finally, stem cell seeding densities has a significant effect on proliferation rate [52, 53]. The slices cultured in growth media displayed a higher metabolic activity likely indicating that the hASCs have higher metabolisms than mature adipocytes. Consistent with histological findings, tissue slices cultured in differentiation media had a higher triglyceride content, where slices cultured in growth media had minimal triglyceride content (Figure 3 G). Overall, these results collectively indicate that the DLM can support hASC differentiation to mature unilocular adipocytes offering a method of creating 3D perfusable adipose tissue models for lean patients.

**Figure 3.**
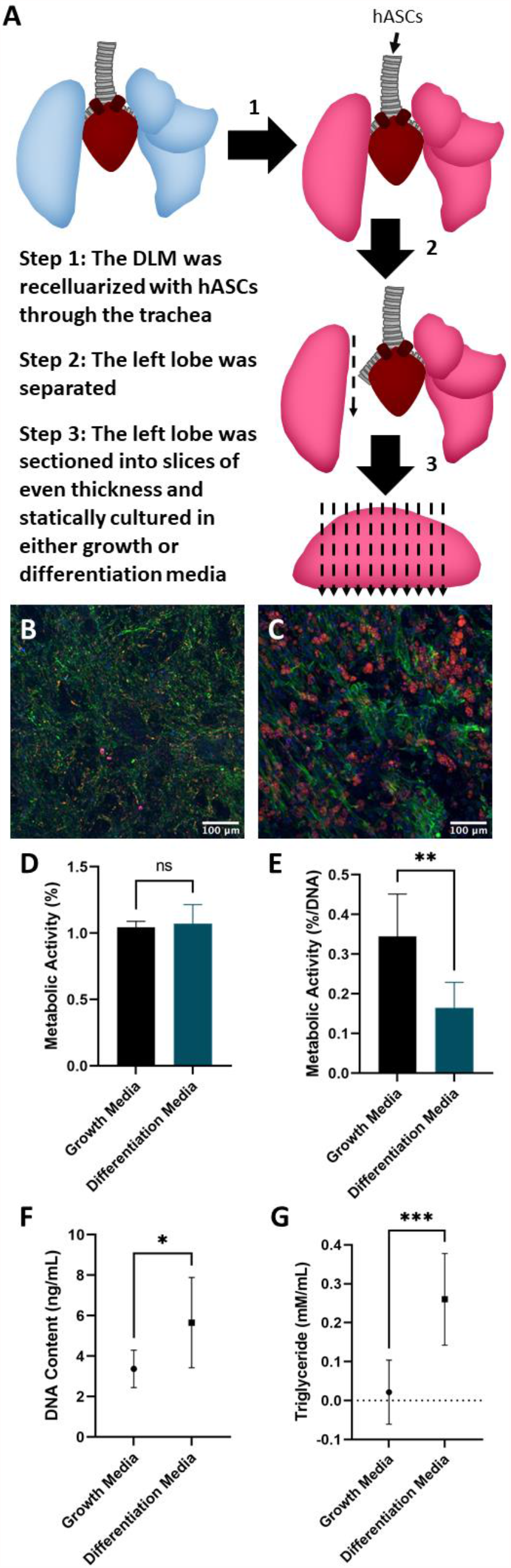
DLM seeded with hASCs cultured in static conditions supported differentiation into mature adipocytes with uniocular lipid droplets. (A) Schematic showing how the hASC seeded DLM was processed. (B) DLM cultured in growth media did not experience significant lipid accumulation in hASCs over the 4-week culture. (C) DLM cultured in differentiation media resulted in a high amount of lipid accumulation with a mixture of multilocular and unilocular adipocytes. (B-C) red – adipocytes, green – actin/native ECM (collagen autofluorescence), blue – nuclei. Scale bars = 100 µm. (D) A resazurin assay was used to measure metabolic activity after 1 week of culture indicating no statistically significant difference between groups (data was not normalized as the same number of cells was seeded into each group). (E) At 4 weeks of culture the resazurin assay indicated that hASCs cultured in the DLM in differentiation media were significantly less metabolically active (results were normalized to the final DNA content to account for proliferation over the culture period). (F) DNA content of the final scaffolds was measured, and the differentiation media resulted in significantly more DNA (cells). (G) Triglyceride content significantly increased in the differentiation group compared to the growth group. This strongly supports the idea that the DLM construct can support hASC differentiation into mature adipocytes. Statistical significance for all data was determined using unpaired t-tests. Asterisks indicate significant (* p<0.05, ** p<0.01, *** p<0.001), while nonsignificant differences are labeled as ns.

### 2.4. Static versus Perfusion Culture

For testing culture conditions, we seeded mature adipocytes directly into the DLM, as this cell source does not require expansion. To investigate if perfusion culture improved outcomes in the DLM over static culture, the left lobe was harvested on Day 0, sliced, and cultured statically while the right lobes were subjected to perfusion culture at a rate of 1.5 mL/min (Figure 4).

**Figure 4.**
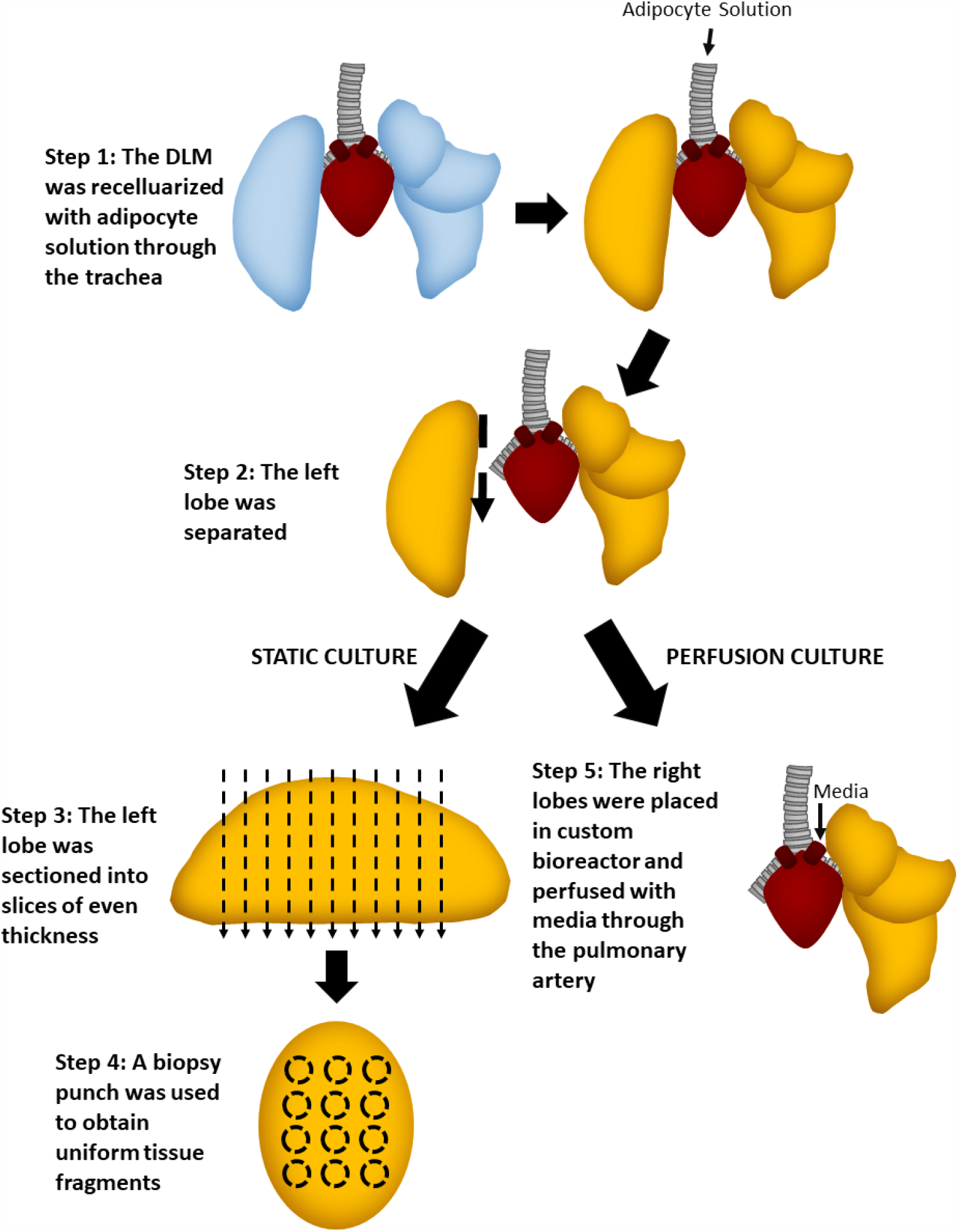
Schematic showing how the lobes were processed to investigate static versus perfusion culture.

Metabolic activity was measured on days 0, 1, 3, and 5 in static (Figure 5 D) and perfusion (Figure 5 E) cultures. In static culture, there was a slight but significant increase in metabolic activity from day 0 to 1. This increase could have been a result of the cells recovering from processing. From day 1 to 3 a steady metabolic activity was detected, after which it significantly decreased from day 3 to 5.

**Figure 5.**
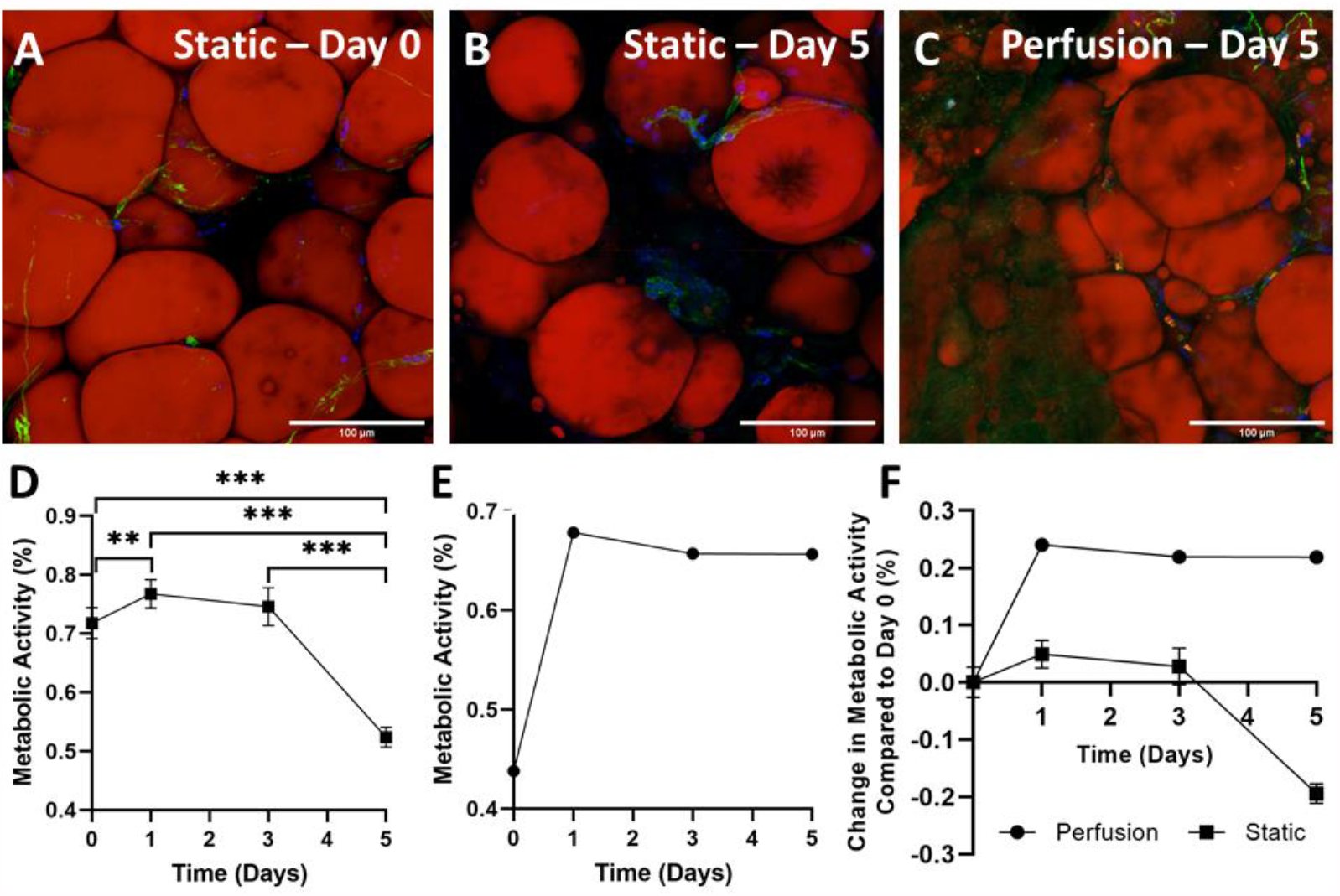
Adipose tissue filled DLMs cultured in perfusion conditions had a more stable metabolic activity compared to static cultures. (A-B) Static samples were imaged on day 0 and day 5 (red – adipocytes, green – actin/native ECM, blue – nuclei). Scale bars = 100 µm. (C) Perfused lobes were imaged at day 5 (red – adipocytes, green – actin/native ECM, blue – nuclei). Scale bars = 100 µm. (D) Resazurin was used to monitor the metabolic activity in static culture conditions, where static culture resulted in a decrease from day 3 to day 5. (E) Resazurin was used to monitor the metabolic activity in perfusion culture. Perfusion culture resulted in a stable metabolic activity over 5 days. (F) The metabolic activity was measured in both culture conditions and is represented as the change in metabolism compared to day 0. Perfusion culture resulted in a more stable metabolic activity over 5 days that did not drop as the static culture did. Statistical significance for metabolic activity data was determined using a one-way ANOVA test followed by a Tukey’s post hoc test. Asterisks indicate significant (** p<0.01, *** p<0.001).

This decline in metabolic activity was not unexpected, as we have seen in other 3D adipogenic perfusion culture systems a trend towards increased lactate dehydrogenase secretion in static over perfused cultures [29]. Furthermore, others have also reported lower metabolic rates in statically cultured cells compared to those in perfusion [54]. Additionally, static samples were fixed and stained on day 0 and day 5 (Figure 5 A-B). At both timepoints adipocytes and stem cells are observed. However, at the 5-day timepoint the adipocyte density is lower with a slight increase in extracellular lipid droplets. Compared to the whole lobes cultured in static conditions, the sliced static culture resulted in less extracellular lipid droplets. This is mostly likely because the lipids could diffuse out due to the increased scaffold to media surface area. Histological findings, paired with the decrease in metabolic activity, indicate that although static culture supports adipocyte and stem cell metabolism short-term, this is not a practical solution long-term.

In comparison to the static culture, the lobes cultured using perfusion techniques demonstrated stable metabolic activity (Figure 5 E). There was a drastic increase in metabolic activity from day 0 to day 1, which could have been due to the cells recovering from the seeding process. Following the increase from day 0 to day 1, the metabolic activity remained constant. Because the lobes were cultured as whole lobes, imaging could only be done at the final timepoint (Figure 5 C). At the 5-day timepoint, both adipocytes and stem cells were observed. The adipocyte density was also similar to the day 0 image of the sliced DLM (Figure 5 A).

Though metabolic activity was measured using the same protocols, differences in experimental setup prevent the static versus perfusion culture results from being directly comparable as the volume of resazurin media used to measure metabolic activity in static culture was significantly less than that used in perfusion culture and the number of cells was also different. Additionally, unlike the cells in static culture that came in direct contact with the resazurin, in perfusion culture the resazurin had to perfuse from the vasculature into the alveolar spaces to be metabolized. To demonstrate the trends from static and perfusion culture on the same plot, the metabolic activity of both cultures was normalized to the day-0 measurement (Figure 5 F). While both static and perfusion groups experienced an increase in metabolic activity from day 0 to day 1, the increase was much larger for the perfused lobes. Both groups experiencing this increase further supports the concept that within the first 24 hours the cells are still recovering from the seeding process. Interestingly, from day 1 to 3 metabolic activities in both groups remained constant. Day 5 is where the biggest difference between the two groups can be seen. While the metabolic activity in the perfusion culture group remained constant, the DLM cultured statically experienced a drastic reduction in metabolic activity. Therefore, the steady metabolic activity and consistent cellular appearance, with minimal extracellular lipids, indicated that perfusion culture offers improved outcomes compared to static culture.

### 2.5. Perfusion Culture of Adipocyte and Endothelial Cell Seeded Lung Matrix

Due to the high lipid-content of large, fragile adipocytes, exposure to high perfusion rates should be avoided [10, 11]. The addition of endothelial cells lining the vascular tree of the DLM offers a unique advantage by creating a physiologically relevant barrier to shield the adipocytes from high shear stresses. To create a 3D vascularized adipose tissue model, the DLM offers easy access for intratracheal cell delivery of adipocytes and large alveolar volume capacity for high-density cell filling, while the pulmonary artery can be used to seed endothelial cells (Figure 6 A). Tile scan images indicated adipocytes were distributed throughout the entire DLM (Figure 6 B), while the endothelial cells repopulated the vasculature tree, surrounding the perimeter of the lung with interior branching evident in the individual tile images (Figure 6 C-D). The endothelial cells can be observed running through areas of heavy adipocyte populations and in some cases the endothelial cells can be seen branching from a larger blood vessel into smaller blood vessels (Figure 6 C).

**Figure 6.**
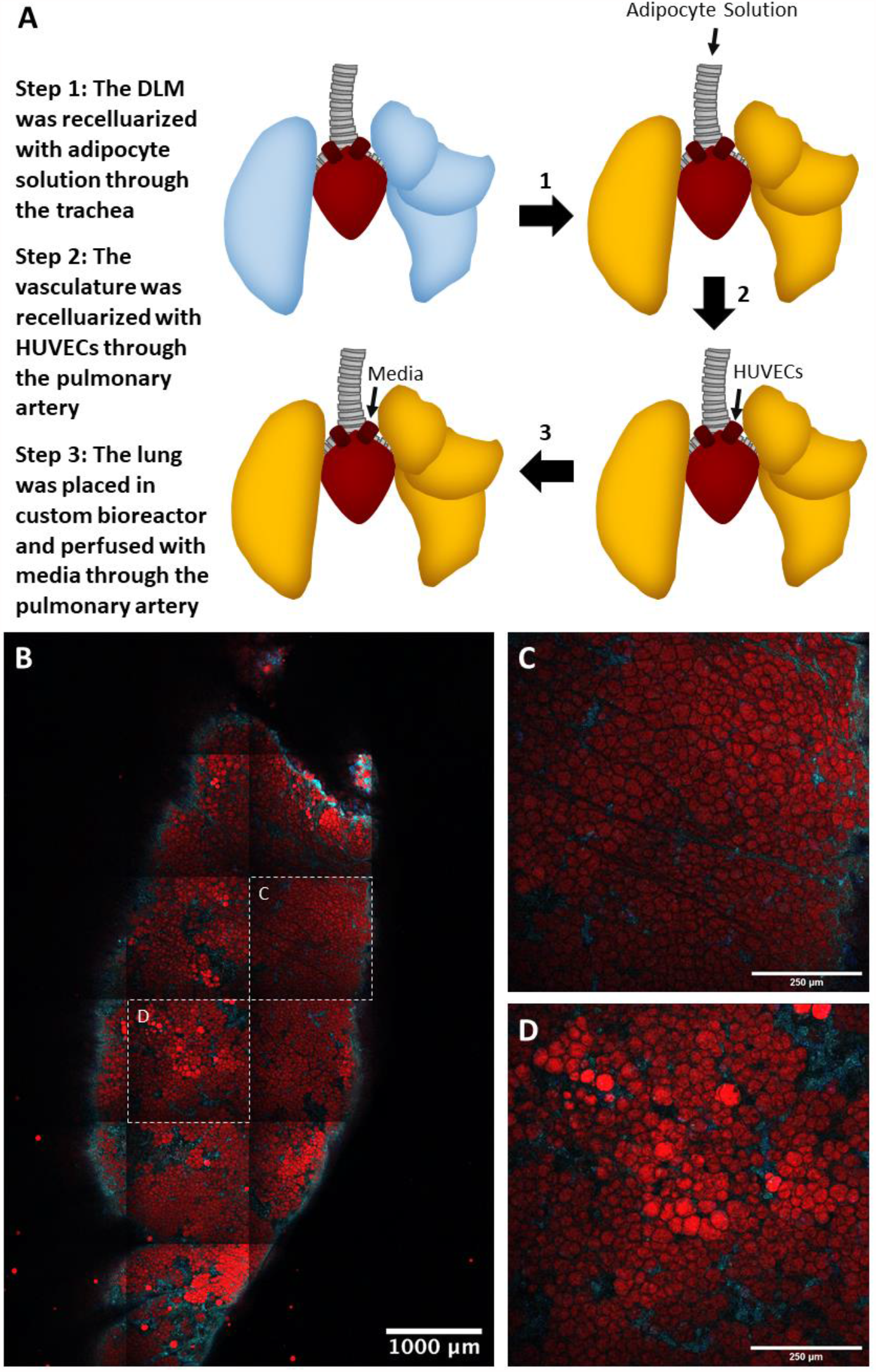
Effective delivery of adipocytes and endothelial cells into the alveolar and vascular compartments of the DLM. (A) Schematic showing how adipocytes and endothelial cells were seeded into the DLM. (B) Tile scan of entire left lobe (red – adipocytes, green – endothelial cells, blue – nuclei). Endothelial and nuclei stains are present in the same areas giving a teal appearance. Image shows that adipocytes seeded through the trachea remain in the inner portion of the lobe and are surrounded by the endothelial cells seeded through the vasculature. Scale bar = 1000 µm. (C-D) Higher magnification images of the tile scan where the scale bars = 250 µm.

### 2.6. Limitations

While this approach is a promising advancement for adipose tissue regenerative medicine there are several limitations in study design. The number of time points was constrained by the number of lung lobes (5). Additionally, differences in lobe size further limited the ability to run parallel studies. For instance, the accessory lobe is considerably smaller than all other lobes. Like any studies using human adipocytes, there are patient to patient differences, such as size, quantity, and metabolic function. To mitigate the differences slightly, adipocytes were isolated from subcutaneous abdominal fat procured from non-diabetic, female patients (ages between 39-55).

## 3. Materials and Methods

### 3.1. Adipose Tissue Culture on Silk Scaffolds

#### 3.1.1. Preparation of Aqueous Silk Fibroin

Silk fibroin solutions were prepared following an established protocol [55]. Whole cocoons (OliverTwistsFibres, Durham, UK) were cut into small pieces and were boiled in a 0.02 M aqueous solution of sodium carbonate (Na2CO3) (Sigma-Aldrich, St. Louis, MO) for 30 minutes to degum the fibroin fibers. The remaining fibers were rinsed and allowed to dry overnight in ambient conditions. The dry fibroin was then dissolved in a 9.3 M aqueous solution of Lithium Bromide (LiBr) (Sigma-Aldrich, St. Louis, MO) at 60°C for 4 hours. This solution was placed into a set of dialysis cassettes (Thermo Fisher Scientific, Waltham, MA) and spun in milli-Q water for 48 hours. The milli-Q water was changed a total of 6 times over the 48 hours. The remaining solution was removed from the cassettes and centrifuged at 4°C and 4800 rpm for 20 minutes. This was repeated to ensure purity.

#### 3.1.2. Preparation of Silk Scaffolds

Silk scaffolds were prepared as described previously [55]. Aqueous silk was lyophilized and then dissolved in a 17% hexafluoroisopropanol solution (Sigma-Aldrich, St. Louis, MO) overnight. The solution was poured over Sodium Chloride (NaCl) (Sigma-Aldrich, St. Louis, MO) crystals with diameters between 500 and 600 μm. The containers were sealed for 24 hours. After the silk permeated through the salt crystals in the container, they were opened and allowed to dry for 24 hours. The dried scaffolds were placed in methanol (PHARMCO-AAPER, Brookfield, CT) for 24 hours to induce β-sheet formation. After methanol annealing, the scaffolds were dried in a chemical hood for 24 hours. The scaffolds were then rinsed for 2-3 days to remove salt from the pores. Finally, the scaffolds were cut into cylinders of 2 mm height and 4 mm diameter.

#### 3.1.3. Adipose Tissue Processing

Adipose tissue was processed using the LIPOGEMS kit (LIPOGEMS International, Norcross, GA). The adipose tissue was first harvested by either using the cannula or manually dissecting the adipose tissue from the skin and blending. For the cannula groups, the adipose tissue was then injected into the LIPOGEMS device and inverted repeatedly causing the enclosed metal ball bearings to evenly break up the tissue into micro-fragments suitable for seeding into the DLM (Figure 1 A). The fragmented tissue was removed from the LIPOGEMS device. Finally, depending on the group, tissues were filtered with a 1 mm secondary filter. This resulted in four experimental groups: Cannula, Cannula+Filter, Blender (alone), Blender+Filter.

#### 3.1.4. Seeding Scaffolds and Culture Conditions

Scaffolds were added to processed adipose tissue and placed in an incubator for 30 minutes. The scaffolds were then transferred to 24 well plates and placed in an incubator for 30 minutes. Media was added after the 30-minute incubation. The scaffolds were cultured in cell culture media (DMEM, 10% fetal bovine serum (FBS), 1% penicillin-streptomycin (pen-strep)) (Thermo Fisher Scientific, Waltham, MA). Media was changed every other day. N=14 for each experimental group.

#### 3.1.5. Live/Dead Staining

Scaffolds from each experimental group were stained with calcein and ethidium (Thermo Fisher Scientific, Waltham, MA). These scaffolds were imaged using a Zeiss LSM 700 Confocal microscope. N=2 for each experimental group.

#### 3.1.6. Imaging

Scaffolds from each experimental group were stained with AdipoRed (Lonza, Basel, Switzerland) for 30 minutes at a 1:35 dilution. These scaffolds were imaged using a Nikon Multiphoton microscope. Collagen was detected using Second Harmonic Generation (SHG). N=2 for each experimental group.

#### 3.1.7. De novo Protein Synthesis

Scaffolds from each experimental group were cultured in media with 50 µM of Ac4GalNAz (Click Chemistry Tools, Scottsdale, AZ) for 3 days (days 4-7 of culture). The scaffolds were then incubated in culture media with 50 µM of DBCO-Cy5 (Sigma-Aldrich, St. Louis, MO). The scaffolds were imaged using the resonant scanner on a Nikon Multiphoton microscope. N=2 for each experimental group.

#### 3.1.8. Picogreen Assay

Picogreen assays (Thermo Fisher Scientific, Waltham, MA) were performed on scaffolds from each experimental group. The assay was performed following the manufacturer’s procedure to assess DNA content. N=6 for each experimental group.

#### 3.1.9. Metabolic Activity Assay

Resazurin (Thermo Fisher Scientific, Waltham, MA) was diluted to 1 mM with phosphate buffered saline (PBS) (pH 7.4). This was diluted further to 0.05 mM solution using the cell culture media. 1 mL of the resazurin solution was placed in each of the 24 wells. The well plate was placed in the incubator for 2 hours. Using a plate reader, the absorbance at 570/600 nm was measured. The metabolism was measured after 4 days of culture. N=8 for each experimental group.

#### 3.1.10. Glycerol Secretion

Media was collected on day 4 and glycerol concentrations were measured using a glycerol assay (BioAssay Systems, Hayward, CA). The assay was performed following the manufacturer’s procedure. N=8 for each experimental group.

#### 3.1.11. Statistics

GraphPad Prism 8.3.0 software was used for all statistical analyses. For this experiment, a one-way ANOVA test followed by a Tukey’s post hoc test was used to determine statistical significance between metabolic activity, DNA content, and glycerol secretions using different processing method (cannula, cannula+filter, blender, blender+filter). For all experiments, significance was defined as *p*<0.05.

### 3.2. Decellularizing the Lung

All studies were approved by the Institutional Animal Care and Use Committee (IACUC) at Carnegie Mellon University (Protocol Number: PROTO201700029). Sprague Dawley rats were euthanized by carbon dioxide overdose. Rat lungs were explanted and decellularized [23, 56]. Briefly, the trachea and the pulmonary artery were cannulated, and the left atrium was lacerated to allow decellularization solutions to easily drain from vasculature. The pulmonary vasculature was perfused via the pulmonary artery cannula using a gravity-driven apparatus suspended approximately 50 cm above the lung. The lungs were perfused with 1 L of 0.1% (w/v) SDS (Sigma) in deionized water (approximately 2 h). Then the lungs were perfused by deionized water (15 min), 1% (v/v) Triton X-100 (Sigma) in PBS (10 min) and followed by three washes with 1 L of PBS to remove residual detergent and cellular debris. Decellularized lungs were stored in PBS supplemented with antimycotic/antibiotic at 4°C until used.

### 3.3 Whole Lobe Static Culture

#### 3.3.1. Seeding the DLM

Recellularization of adipocytes was performed in custom-made bioreactors [57] allowing adipocyte delivery through the tracheal outlet and perfusion through the pulmonary artery. Prior to seeding, decellularized lung scaffolds were primed by perfusing 100 mL of adipocyte culture medium (DMEM high glucose, 10% FBS, 1% pen-strep) for at least 1 h (1 mL/min). Next, the cellular solution (adipocytes and SVF suspended in PBS) was injected, where adipocytes were delivered into the alveolar space through the tracheal outlet. Adipose tissue was processed as described in *3.1.3* with a cannula, a LIPOGEMS kit, and a 1 mm secondary filter. A total of 10 mL of adipocytes were infused into the tracheal outlet at 1 mL/min using a syringe pump. The seeded lung scaffold was placed statically in cell culture media in the incubator for 1 hour. For this experiment, the lobes were then separated, and each lobe was cultured statically in a 6 well plate in cell culture media (DMEM high glucose, 10% FBS, 1% pen-strep) (Thermo Fisher Scientific, Waltham, MA). Media was changed on days 1, 3, and 7. The lobes were fixed at different timepoints (Day 0, 1, 3, and 7).

#### 3.3.2. Imaging

Lobes were sliced and stained with AdipoRed, phalloidin (1:200), and DAPI (1:1000). The lobes were imaged using a Nikon Multiphoton microscope.

#### 3.3.3. Glycerol Secretion

Media was collected on day 1, 3, and 7 from the left lobe. The glycerol concentrations were measured using a glycerol assay (BioAssay Systems, Hayward, CA). The assay was performed following the manufacturer’s procedure. N=1 for each timepoint.

#### 3.3.4. Statistics

For this set of experiments, a linear regression was performed on the glycerol concentration data. The R^2^ value was calculated.

### 3.4 Static Culture of sectioned DLM

#### 3.4.1. Seeding DLM

The DLM was seeded as describe in *3.3.1*. Following the 1-hour incubation after seeding, for this set of experiments, the left lobe was then removed and cut into slices (approximately 1 mm thickness). Using a 4 mm biopsy punch, uniform sections of the lobe were isolated. The sections were cultured statically in a 24 well plate and incubated for and additional hour before adding cell culture media (DMEM, 10% FBS, 1% pen-strep) (Thermo Fisher Scientific, Waltham, MA) for 5 days. Media was changed on days 1, 3, and 5.

#### 3.4.2. Imaging

Lobe sections were stained with AdipoRed, phalloidin, and DAPI as described previously. The lobes were imaged using a Zeiss LSM 700 Confocal microscope.

#### 3.4.3. Metabolic Activity

Resazurin (Thermo Fisher Scientific, Waltham, MA) was diluted to 1 mM with phosphate buffered saline (PBS) (pH 7.4). This was diluted further to 0.05 mM solution using the cell culture media. 1 mL of the resazurin solution was placed in each of the 24 wells. The well plate was placed in the incubator for 4 hours. Using a plate reader, the absorbance at 570/600 nm was measured. The metabolism was measured on days 0, 1, 3, and 5. N=16 for day 0. N=8 for days 1, 3, and 5.

#### 3.4.4. Statistics

this experiment, a one-way ANOVA test followed by a Tukey’s post hoc test was used to determine statistical significance between metabolic activity on different days.

### 3.5. Perfusion Culture with Whole Adipose Tissue

#### 3.5.1. Seeding DLM

The DLM was seeded as describe in *3.3.1*. Following the 1-hour static incubation after seeding, for this set of experiments, the left lobe was removed for static culture, while the right was cultured using the perfusion bioreactor. The right lobe was placed statically in the bioreactor with media for 2 hours.

#### 3.5.2. Perfusion Culture

After 2 h of static culture to allow cell attachment, perfusion was initiated at 1.5 mL/min through pulmonary artery. 50 mL of adipocyte culture medium (DMEM high glucose, 10% FBS, 1% pen-strep) was used. On Day 2, 50 mL of the culture medium was removed and replaced with 50 mL fresh medium. A full medium change was performed on Day 4 and the recellularized lung scaffold was harvested on Day 7 for further analysis.

#### 3.5.3. Imaging

Lobes sections were stained with AdipoRed, phalloidin, and DAPI as described previously. The lobes were imaged using a Zeiss LSM 700 Confocal microscope.

#### 3.5.4. Metabolic Activity

The Resazurin assay was performed as described in 3.4.4. on days 0, 1, 3, and 5.

### 3.6. Static Stem Cell Seeded DLM Slices

#### 3.6.1. Isolation of human adipose derived stem cells (hASCs)

hASCs were isolated from subcutaneous adipose tissue. The cells were isolated by mechanically blending the adipose tissue and incubating it in a collagenase solution (0.1% collagenase, 1% bovine serum albumin, 98.9% phosphate buffer solution) (Thermo Fisher Scientific, Waltham, MA) at a 1:1 ratio. The mixture was placed in a cell culture incubator for 1 hour and then centrifuged to isolate the stromal vascular fraction. The cells were resuspended in media (DMEM with 10% FBS and 1% pen-strep), centrifuged, and seeded into flasks.

#### 3.6.2. Seeding DLM

hASCs were lifted from cell culture flasks using 0.25% trypsin-EDTA. A total of 50 million hASCs (5 million cells/mL) were infused through the trachea to repopulate the acellular rat lung scaffold at 1 mL/min using a syringe pump. After overnight culture in custom-made bioreactor, the lung scaffold was infused with 3 mL of 2 % (w/v) low melting temperature agarose through trachea and allowed to gel for 5 min at RT. The hASC-seeded lung did not swell as much as the adipocyte-seeded lung, so low melting temperature agarose was added to facilitate accurate lobe dissection. Using a sterile blade, the lung lobes were sliced to a 1 mm thickness. Sliced lung scaffolds were placed in a 24-well plate and cultured statically in growth media (DMEM/F12, 10% FBS, 1% pen-strep) or differentiation media (DMEM/F12, 10% FBS, 1% pen-strep, 1μM insulin, 0.5 mM IBMX, 0.05 mM indomethacin, and 1 μM dexamethasone). N=8 for growth media and N=9 for differentiation media.

#### 3.6.3. Imaging

Slices were stained with AdipoRed, phalloidin, and DAPI as described previously. The slices were imaged using a Zeiss LSM 700 Confocal microscope.

#### 3.6.4. Metabolic Activity

The Resazurin assay was performed as described in *2.4.4*. at 1 week and 4 weeks. N=8 for growth media and N=9 for differentiation media. The results from week 1 were not normalized as the same number of cells were in each slice due to the seeding and slicing procedure.

#### 3.6.5. Picogreen Assay

The picogreen assay (Thermo Fisher Scientific, Waltham, MA) was performed on 7 scaffolds for each group. The assay was performed following the manufacturer’s procedure to assess DNA content. The data was also used to normalize the metabolic assay results for week 4. N=7 for growth media and differentiation media.

#### 3.6.6. Triglyceride Concentration

Lysed cell solution was used to determine triglyceride content using a triglyceride assay (BioAssay Systems, Hayward, CA). The assay was performed following the manufacturer’s procedure. N=7 for each experimental group.

#### 3.6.7. Statistics

To compare metabolic activity, DNA concentration, and triglyceride concentration when cultured in growth versus differentiation media, unpaired *t* tests were used.

### 3.7. Vascularized Whole Lobe Perfusion Culture

#### 3.7.1. Adipose Tissue Processing

To further liquify the solution and ensure that only adipocytes were seeded and penetrated in the DLM, following mechanical disruption adipose tissue was incubated in a collagenase solution (0.1% collagenase, 1% bovine serum albumin, 98.9% phosphate buffer solution) (Thermo Fisher Scientific, Waltham, MA) at a 1:1 ratio. The mixture was placed in a cell culture incubator for 4 hours and then centrifuged. The oil layer was removed using an aspirator and the adipocyte layer was then filtered using a 355 μm sieve to remove any remaining pieces of ECM that could clog the cellular injection into the decellularized lung.

#### 3.7.2. Endothelial Cell Culture

Human umbilical vein endothelial cells (HUVECs, Lonza) were maintained in 0.1% (w/v) gelatin-coated flasks with complete EGM-2 (Lonza) supplemented with 1% penicillin-streptomycin (Gibco). The cells were passaged when reaching 80-90% confluency. Cells at passage 4 were used for all the experiments.

#### 3.7.3. Seeding DLM

40 million HUVECs were seed into the DLM through the existing pulmonary vasculature. Two hours after HUVECs were seeded, 10 mL of adipocyte cell solution were seeded into the DLM through the tracheal outlet. Seeded DLM was then cultured using perfusion speeds of 1.5 mL/min and adipocyte maintenance media (DMEM low-glucose, 10% FBS, 1% pen-strep) (Thermo Fisher Scientific, Waltham, MA) cell culture media. On Day 2, 50 mL of the culture medium was removed and replaced with 50 mL fresh medium. A full medium change was performed on Day 4 and the recellularized lung was harvested on Day 7 for further analysis.

#### 3.7.4. Imaging

The DLM was sliced and stained with AdipoRed, VE-cadherin, and DAPI. The slices were imaged using a Zeiss LSM 700 Confocal microscope.

## 4. Conclusion

Overall, we found that the DLM supported adipocyte viability, function, and differentiation in a 3D matrix. Both adipocytes and hASCs were successfully delivered and remained in the alveolar space even after weeks of culture. When seeded into the DLM, the adipose derived cells maintained their morphology and functionality in both static and perfusion cultures. However, perfusion culture offered enhanced outcomes over static culture due to the lower extracellular lipid accumulation and stable metabolic activity. Furthermore, the DLM successfully promoted adipogenic differentiation of hASCs which is critical for lean patients. The incorporation of endothelial cells in the vasculature is essential to create a fully functional vascularized adipose tissue graft. The endothelial cells seamlessly integrated into the vascular tree of the DLM. These results support that the seeded DLM could be a promising advancement in vascularized adipose tissue grafts for large defect filling.

